# Wing Metric Variation in *Aedes aegypti* Effect of Altitude on Wing Metric Variation of *Aedes aegypti* (Diptera: Culicidae) in a Region of the Colombian Central Andes

**DOI:** 10.1101/2020.01.29.924746

**Authors:** Luis M. Leyton-Ramos, Oscar Alexander Aguirre-Obando, Jonny Edward Duque, Víctor Hugo García-Merchán

## Abstract

In mosquitoes of medical importance, wing shape and size can vary with altitude, an aspect that can influence dispersion and, consequently, their vector capacity. Using geometric morphometry analysis, *Aedes aegypti* wing size and shape variation of males and females was studied in four altitudes in the second-smallest department in Colombia: 1.200 m (Tebaida), 1.400 m (Armenia), 1.500 m (Calarcá), and 1.700 m (Filandia). Wing shape in males (P < 0.001) and females (P < 0.001) was significantly different through the altitudinal gradient; in turn, wing size in males followed the altitudinal gradient (Males R^2^ = 0.04946, P = 0.0002), Females (R^2^ = 0.0011, P = 0.46). Wing allometry for males (P < 0.001) and females (P < 0.001) was significant. Likewise, the shape and size of the wings of males (P < 0.001) and females (P < 0.001) had significant fluctuating asymmetry. It is concluded that, in a small scale with an altitudinal variation of 500 meters, it is detected that the size and shape of the wings varied in *A. aegypti*, principal vector of dengue, chikungunya, and Zika. The fluctuating asymmetry is present in the individuals studied and could be associated with environmental effects caused by vector control campaigns present in some sampling locations.

## 1. Introduction

*Aedes (Stegomyia) aegypti* (Linnaeus, 1762) is an urban anthropophilic mosquito from Africa, distributed in the world’s tropical and sub-tropical regions [1]. In the Americas, this mosquito is present in almost every country, considered the principal vector of dengue (DENV), Zika fever (ZIKV), and chikungunya (CHIKV) [2–4]. In Colombia, *A. aegypti* is registered in 80% of the country up to 2.300 m [5]. Nevertheless, still unknown is the epidemiological impact altitude exerts on the population dynamics of *A. aegypti* in areas, like the Andean region. It has been observed that the altitude in the zones where the mosquito inhabits has a direct impact on the abundance, geographic distribution, vector capacity, epidemiology, and pathogenicity of the mosquitoes [6]. Additionally, in culicids, the range of altitudinal distribution may be modified by increased global temperature [7], a phenomenon observed in the natural populations of *A. aegypti* of the Americas, including Colombia [5].

In *A. aegypti*, the size of the individuals has been associated with components of the reproductive success [8, 9] [9]. Bigger *A. aegypti* individuals (*per se*, bigger wingspan) could be more involved in the transmission of arthropod-borne virus (arbovirus), like dengue, than smaller ones [10]. In addition, bigger individuals have been associated with a higher frequency of feeding from blood in human hosts [11], greater survival, and fertility [12]. On the contrary, smaller mosquitoes (hence, with smaller wingspan) may have a higher number of feeding events throughout their lives, which can increase infection levels and arbovirus dissemination [9, 13, 14]. Furthermore, the biological shape is an outstanding aspect of the phenotype of an organism and provides a link between the genotype and the environment [15]. Said shape has been studied in insects, like butterflies and fruit flies, with emphasis on the wings, which are a trait associated with load capacity and dispersion [16, 17]. In mosquitoes, wing shape is associated with dispersion capacity [18]. Additionally, wing flapping produces vibrations that generate sounds [19, 20], which are different and are related with the precopulatory behavior [21].

In mosquitoes, it has been noted that temperature (climatic variable inversely proportional to altitude) causes changes in the life cycle, affecting the body size and shape [22]. An inverse relationship has been observed in some cases between temperature and the duration of development [23]. Consequently, at lower temperatures, the transmission of arbovirus may – in some cases – be impeded [24]. Hence, knowing how wing shape and size vary in *A. aegypti*, with relation to altitude, could contribute useful information for its vector control.

The geometric morphometry permits detecting information patterns on the type, ecological relationships, and environmental factors associated to populations present in the area [25–27]. In *A. aegypti*, morphometric analyses on wings have been widely studied to investigate heterogeneity and structuring in natural populations [28–31]. However, very few prior studies have related altitude and wing metric variation in mosquitoes. Studies in northeastern Turkey on *Aedes vexans* between 808 and 1620 m and on *Culex theileri* between 808 and 2.130 m showed variation in wing size and shape. Besides, in *Culex theileri*, a positive correlation was observed between wing size and altitude [18, 32]. Recently, in *Aedes albopictus* of Albania, the region where this Asian mosquito was first registered in Europe, it was observed between 154 and 1559 m shape, size and sex variations among altitudinal populations of these species [33]. Nevertheless, in Colombia and the rest of the world, the metric variation of *A. aegypti* and its relation with the altitudinal gradient has not been studied much [34]. In Colombia, the department of Quindío is on the central mountain range of the Andes and it is the second smallest in geographic extension with altitudes in its urban area ranging from 1.200 to 1.917 m in 1,961 km^2^ [35–37]. Due to the aforementioned, this study sought to explore the wing size and shape variation of *A. aegypti* males and females from an altitudinal gradient of the central mountain range of the Colombian Andes.

## 2. Methodology

### 2.1 Field work

The department of Quindío is the second smallest regarding land area in Colombia, with an extension of 1.961 km^2^, with an altitude range in the urban area from 1.200 (Tebaida) to 1,917 m (Filandia) [35, 36]. Here, for six months, between August (2017) and February (2018), adult individuals of *A. aegypti* were collected in four altitudes of the urban zone. The sampling sites were the following: 1.200 m (Tebaida), 1.400 m (Armenia), 1.500 m (Calarcá), and 1.700 m (Filandia). In Filandia, one of the municipalities with the highest altitude urban settlement in the department, no mosquitoes were found at altitudes above 1.700 m. To select the sampling sites by altitude, 10 points were randomly selected. Each point corresponded to a neighborhood and in each neighborhood, homes were visited where their dwellers permitted. Each home was visited four times per month and each visit lasted from 45 to 60 minutes. In each home sampled, an informed consent was delivered on the objectives of the research and the authorization to conduct the sampling.

The adults were collected through mechanical aspiration through an electric aspirator. After collection, the individuals were sedated and sacrificed with acetone. All collections were made under the framework permit from the Corporación Autónoma Regional del Quindío (CRQ) N° 240 issued for the department of Quindío, Colombia. Thereafter, the specimens were identified at species level by using the dichotomous keys by [38] and [39]. Fig 1 shows the location, as well as the total number of mosquitoes by altitude and sex used in this work.

**Figure 1.**
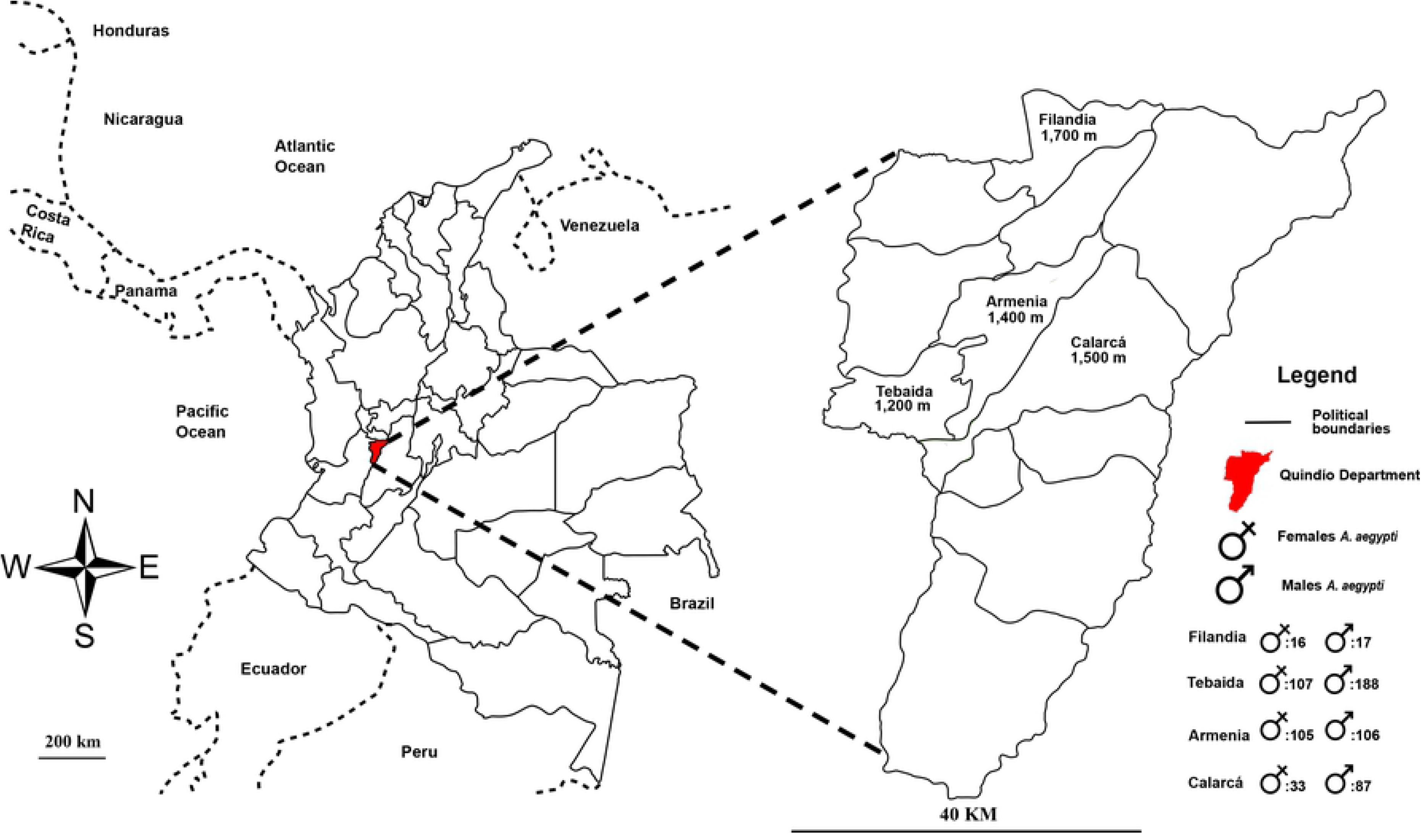
Map with sampling altitudes and total number of males and females of *A. aegypti* by altitude.

### 2.3 Laboratory work

The left and right wings were removed from each male and female mosquito collected; removal was from the base, following the protocol described in [40] and [41]. Each wing was submerged in NaClO solution at 5% to remove scales and rinse them. Thereafter, each wing was submerged in ethanol solution at 99.5% to remove excess NaClO, to be mounted on a slide with ethanol at 70%. The photographs were taken on a stereomicroscope (Zeiss Stemi DV4) with integrated camera (Canon EOS REBEL T3i) with 32X magnification, according to specifications for taking landmarks (LM) for two dimensions (2D) [42].

Each wing was marked with 22 LM [43] type I (Fig 2A), according to the method by Rohlf & Slice [26, 44]. The photos were organized randomly with the tpsUtil software version 7.0 to reduce LM marking bias [45]. All the LM of the wings were marked by using the tpsDig2 program [46]. The LM were located by the same operator to reduce human error in taking points. To guarantee the reproducibility of the experiment, the marking of the LM was carried out twice. To detect atypical LM, the MorphoJ program was used [47].

**Figure 2.**
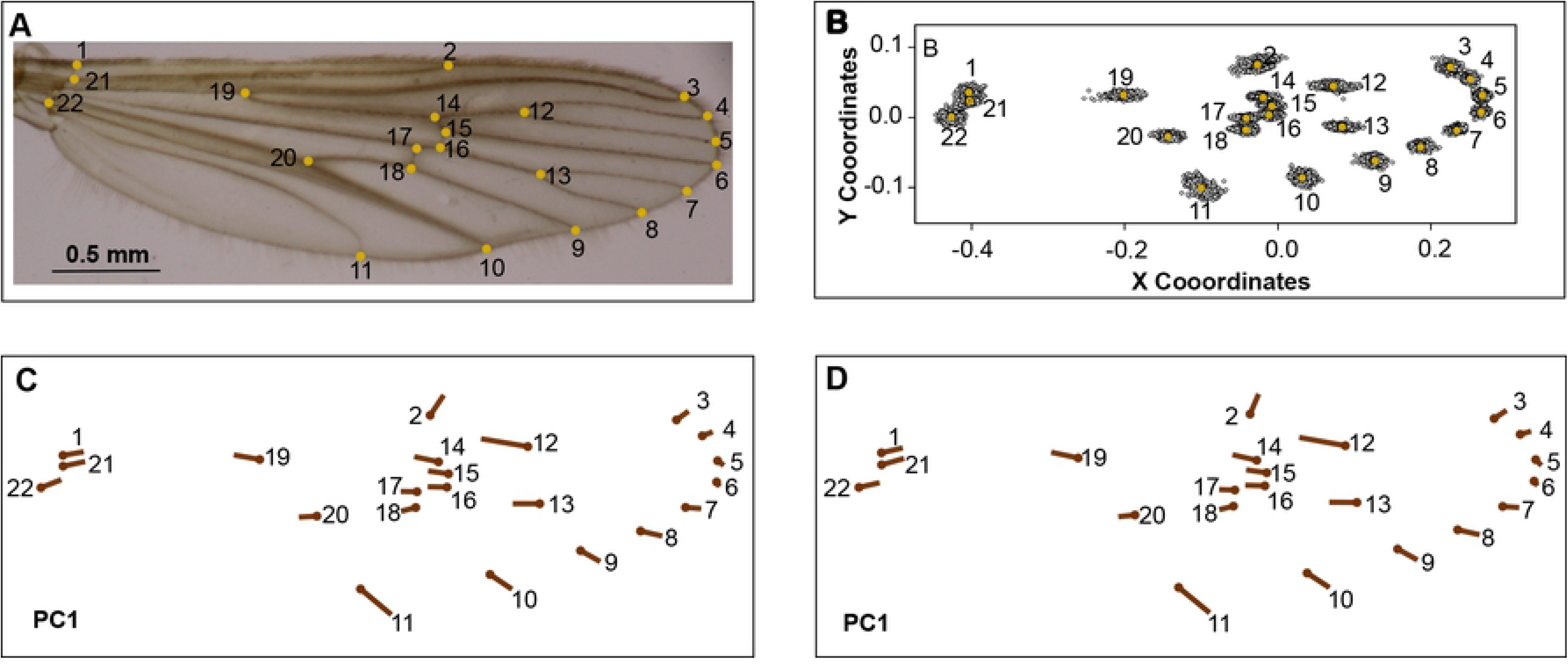
Location of LM, general analysis of Procrustes, and comparison among replicas. (A). Anatomical frameworks used. (B) Procrustes coordinates after the analysis of atypical data. (C) PC1 set of original data. (D) PC1 set of replica data.

### 2.4 Data analysis

From the matrix of data with the coordinates of each LM, the effect of the scale, translation, and rotation was eliminated through a general Procrustes analysis [44, 48]. Thereafter, the Procrustes coordinates were obtained as representative variable of the wing shape and centroid size (CS) of the wing size, which were used in all the analyses performed. To guarantee reproducibility of the data used, the measurement error rate was estimated according to [49]. To visualize the differences of each LM among the original data and their replica, from the principal components obtained from Procrustes coordinates, deformation grids were used [50].

Information from the right wing was used to analyze the shape and size variation in the altitudinal gradient for both sexes. The CS for each sex and altitude was evaluated by using the Kruskal-Wallis non-parametric test. When differences were significant, a pair-wise Mann–Whitney U-test was used, and it was visualized by using a box diagram and a chart to indicate the significant differences among the comparisons. To determine the relationship between size and the altitudinal gradient, a simple linear regression was performed. The allometric influence of wing size within the shape was analyzed through a multivariate regression of Procrustes coordinates [51] in function of the CS, using a permutations test with 10,000 randomizations [52].

The wing shape variation patterns for each sex were visualized through an analysis of principal components analysis (PCA). For each sex, wing shape and its variation in the altitudinal gradient, a Canonical Variable Analysis (CVA) was conducted. In addition, both the PCA and the CVA used deformation grids to determine in what LM did the wing shape variation originate. The shape variation in function of the altitudinal gradient for each sex was evaluated through an analysis of variance (ANOVA) with 10,000 randomizations [53].

With the mosquitoes (Males = 223 and Females = 608) that had both wings in good condition, symmetry or asymmetry were determined. For this, the study used values of the CS and the Procrustes coordinates of the original data and their replica, which were analyzed through a Procrustes ANOVA with 10.000 iterations [54, 55]. All the analyses were performed in the R programming software, version 3.5.0 [56], using the Geomorph package, version 3.1.2 [57], except for the CVA, Mahalanobis distances, and the allometric regression, which were carried out in the MorphoJ program, version 2.6 [47].

## 3. Results

Fig 2B shows that the distribution of the LM used is located continuously. Fig 2C and D indicate that the set of original data and their replica had no variations by the user to mark the 22 LM (user measurement error rate: 1.92%). The CS of the wings of males and females had significant differences (Gl = 1, P < 0.05), being higher in females (4.19 ± 0.38) than in males (3.29 ± 0.30); (Fig.3). In turn, the CS variation in function of the altitudinal gradient for wings from males (Gl = 3, P < 0.05) and females (Gl = 3, P < 0.05) was statistically significant, indicating the CS for males, based on the Mann– Whitney test, statistical differences between the altitudes from 1.400 to 1.500 m, 1.200 – 1.400 m, and 1.200 – 1.700 m. The CS of the wings for females was different between the altitudes 1.400 – 1.500 m and 1.200 – 1.400 m Table. 1. Fig 4A suggests that wings of males have a slight tendency to being bigger at higher altitudes, nevertheless, other variables associated with the mosquito’s development must be analyzed to have better resolution (R^2^ = 0.04946, P = 0.0002), a pattern not observed for wings of females (R^2^ = 0.0011, P = 0.46; (Fig 4B)).

**Figure 3.**
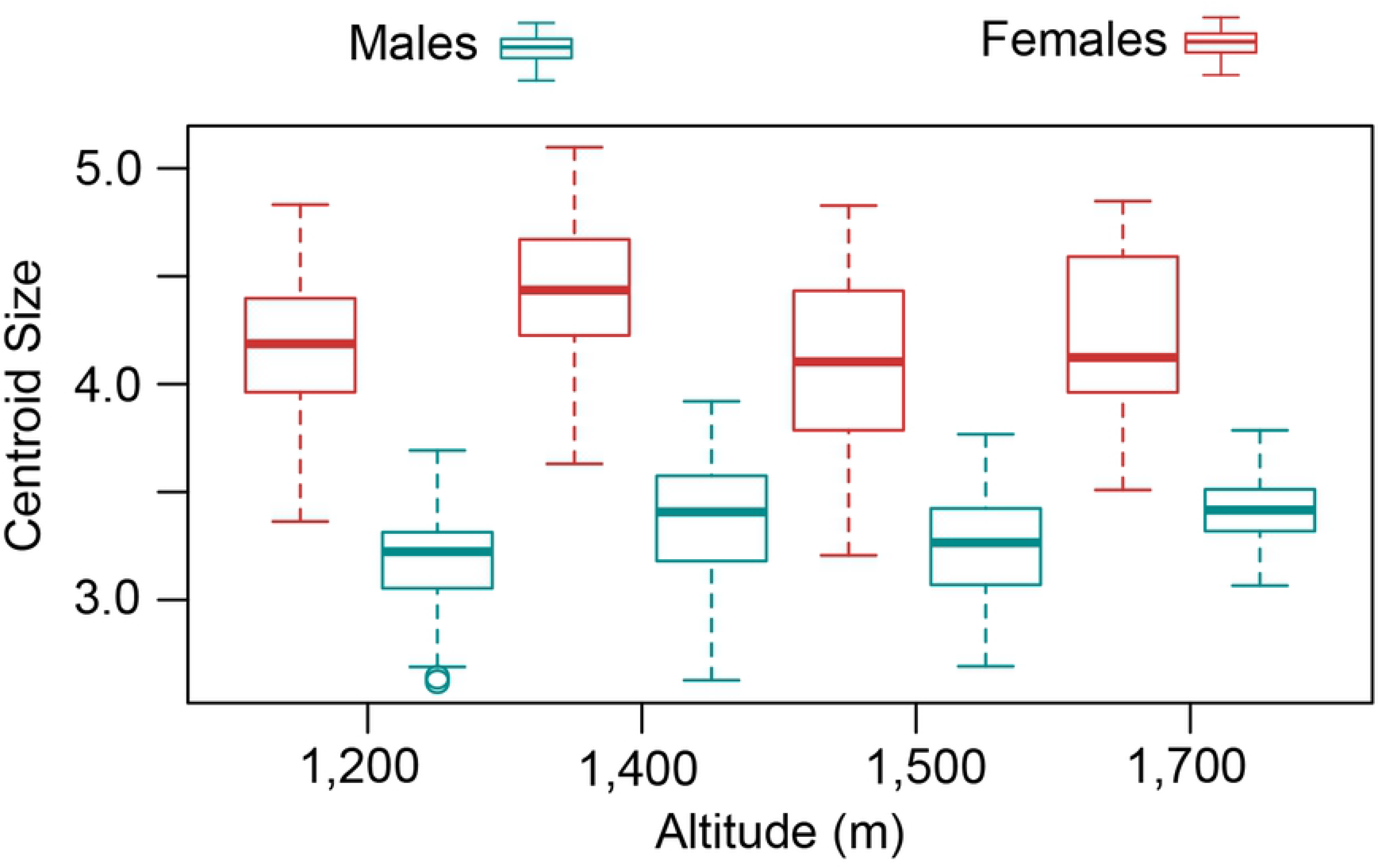
Box diagram for the centroid size and altitude for wings from both sexes of *A*. *aegypti*.

**Table 1.** Significant differences for size by sex in the altitudinal gradient, according to the Pairwise Mann–Whitney U-tests.

| Sex | 1.200 m | 1.400 m | 1.500 m | 1.700 m |
| --- | --- | --- | --- | --- |
| **Females** | a | b | a | b |
| **Males** | a | b | a | ab |

\*Different letters indicate significant differences ( $p < 0.05$ ).

**Figure 4.**
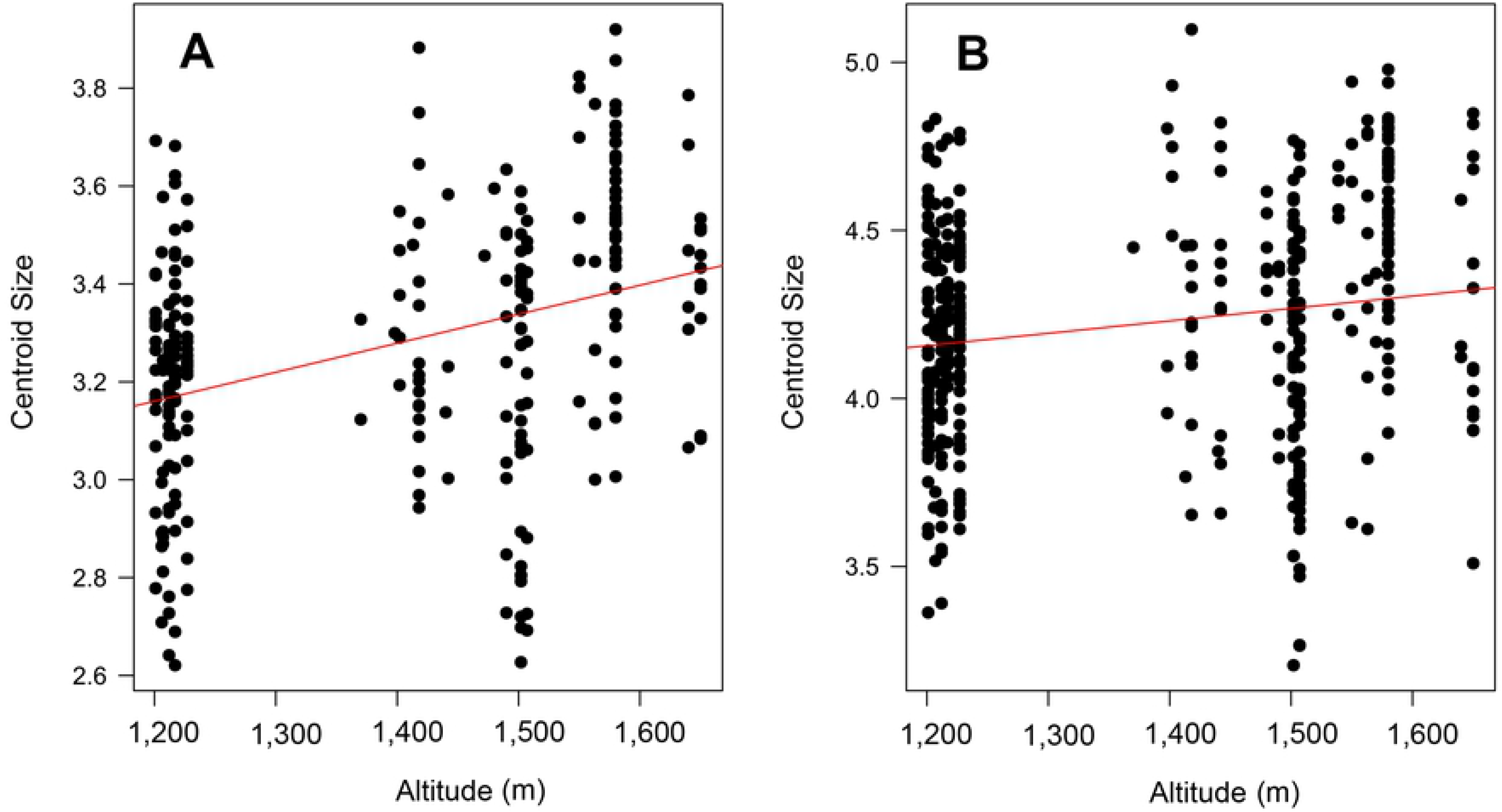
Linear regression for centroid size and altitude for both sexes of *A*. *aegypti*. (A) Males. (B) Females.

The allometry test indicated that the contribution of CS wing shape variation was significant for both sexes (Males P < 0.001, Females P < 0.001), where the percentage of wing shape variance explained by the size was 4.9% for males and 2.5% for females.

The PCA in total explained 53.1% of data variation (PC1 = 43.8%, PC2 = 9.3%). The PC1 separated two groupings corresponding to each sex. The wing shape of males was present in the negative part of PC1 and that of the females in the positive part. In males, wing shape variation was observed for LM from the wing contour (LM: 1-11, 21, 22) in negative sense to PC1 and internal LM (LM 12-20) in positive sense to PC1. For females, the LM from the wing contour were present in positive sense to PC1 and in contrary sense, the internal LM (Fig. 5).

**Figure 5.**
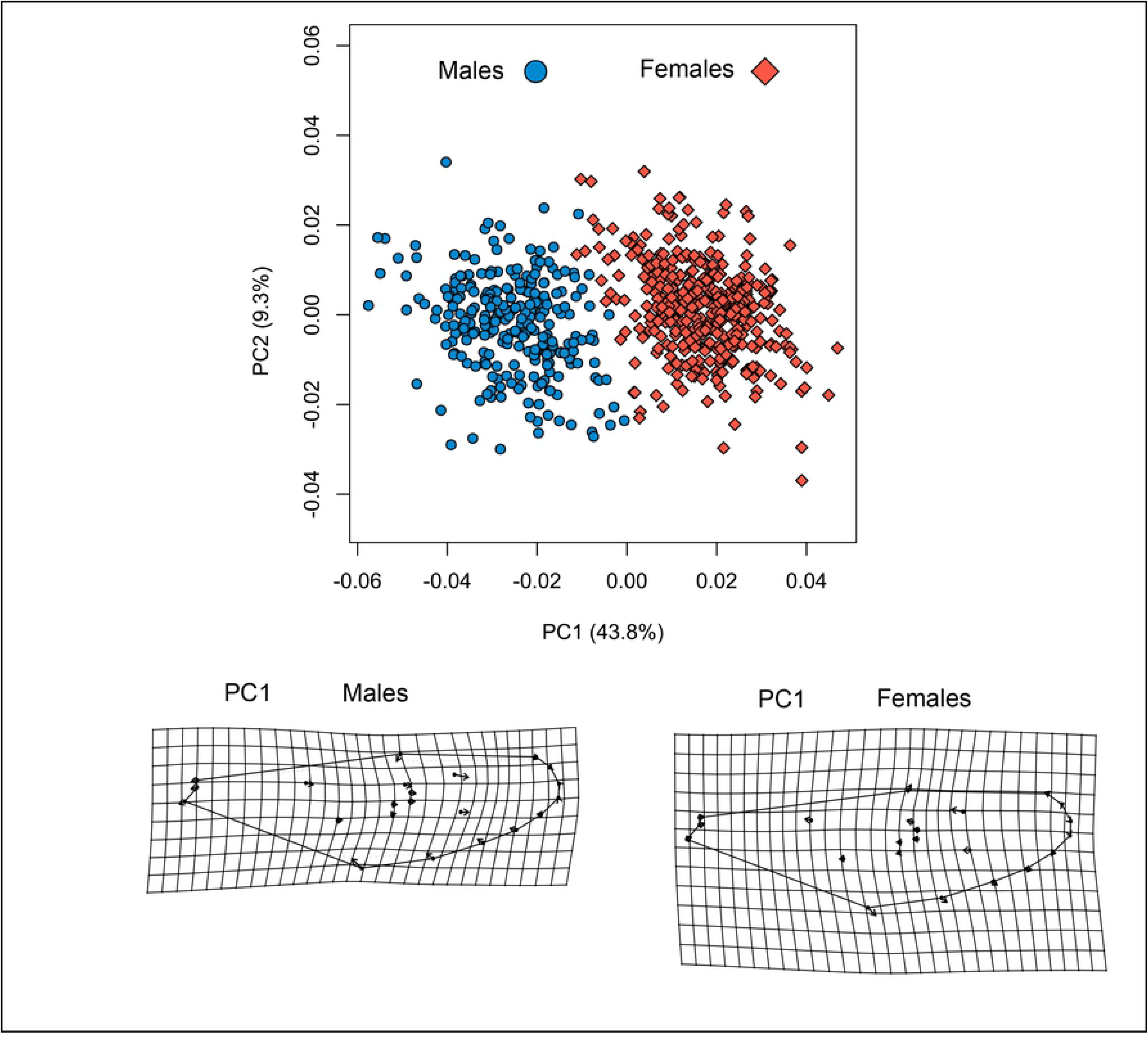
PCA and deformation grid for wing shape between both sexes of *A*.*aegypti*.

The CVA for each sex showed differences in wing shape among mosquitoes located at different altitudes. In males, CV1 and 2 explain – in total – 85.3% of the wing shape variation (CV1 = 62.2%, CV2 = 21%), while in females, this explained 82.9% of the wing shape variation (CV1 = 60.4%, CV2 = 22.4%). These differences in wing shape were supported by the permutations test for distances by Mahalanobis for males (P < 0.001) and females (P < 0.001) (Fig.6). Wing shape in function of the altitudinal gradient for males (F = 3.8251; Gl = 3; P = 0.001) and females (F = 3.5457; Gl =3 P = 0.001) were significant Table 2.

**Figure 6:**
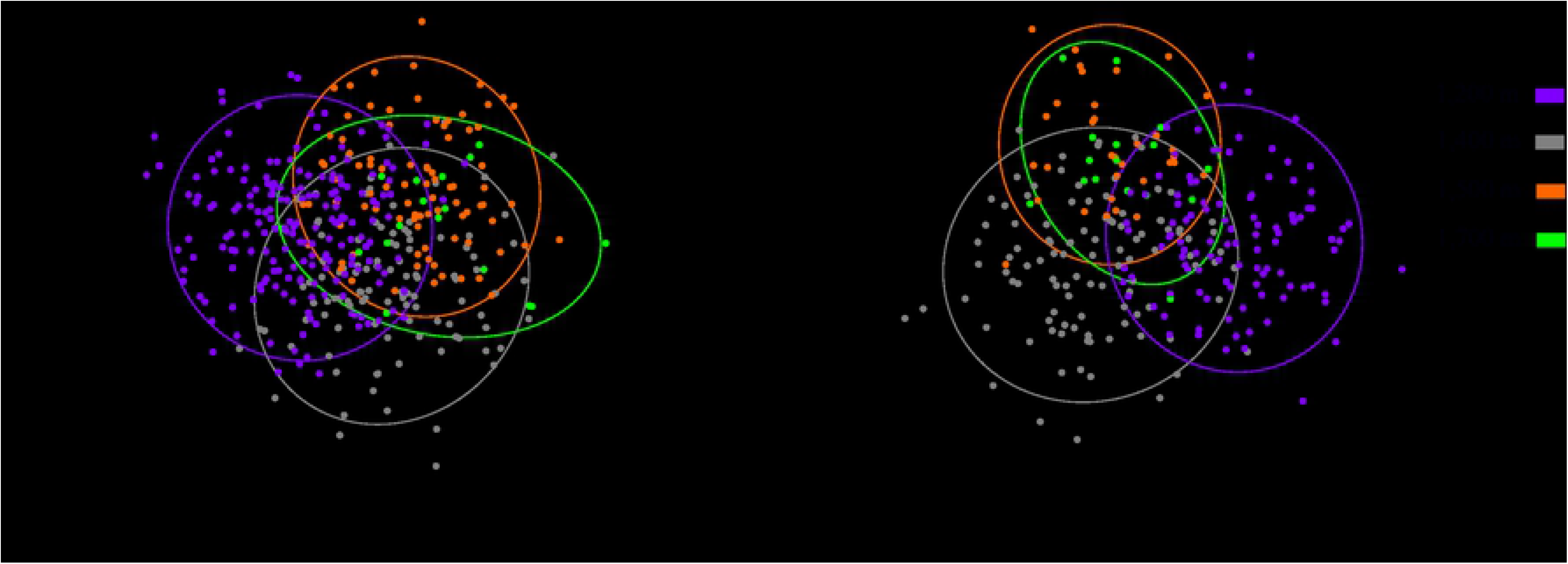
CVA for wing shape in the altitudinal gradient: (A) Females. (B) Males.

**Table 2.** MANOVA of Procrustes for wing shape in function of the altitudinal gradient for males and females of *A. aegypti*.

|  |  | Df | SS | MS | Rsqr | F | Z | p-value |
| --- | --- | --- | --- | --- | --- | --- | --- | --- |
| **Females** | **Altitude** | 3 | 0.007739 | 0.0025796 | 0.02629 | 3.5457 | 5.2723 | 0.0001 |
|  | **Residuals** | 394 | 0.286646 | 0.0007275 | 0.97371 |  |  |  |
|  | **Total** | 397 | 0.294385 |  |  |  |  |  |
| **Males** | **Altitude** | 3 | 0.009286 | 0.0030954 | 0.04274 | 3.8251 | 5.4866 | 0.0001 |
|  | **Residuals** | 257 | 0.207972 | 0.0008092 | 0.95726 |  |  |  |
|  | **Total** | 260 | 0.217259 |  |  |  |  |  |

\* Significant differences ( $p < 0.05$ ).

The bilateral symmetry test for shape indicated no significant variation between the left and right sides for males (Side P = 0.157) and females (Side P = 0.157) of *A. aegypti*. On the contrary, the variation among individuals and its interaction with the wing side, was indeed significant for males (Individuals P = 0.001, Side*Individual P = 0.001; Table 3 and females (Individuals P = 0.0001, Side*Individual P = 0.001; Table 4. For the CS, the bilateral symmetry test indicated variation among individuals, between sides and its interaction for males (Side P = 0.0001, Individuals P = 0.0001, Side*Individual P = 0.001) and females (Side P = 0.0001, Individuals P = 0.0001, Side*Individual P = 0.001).

**Table 3.**
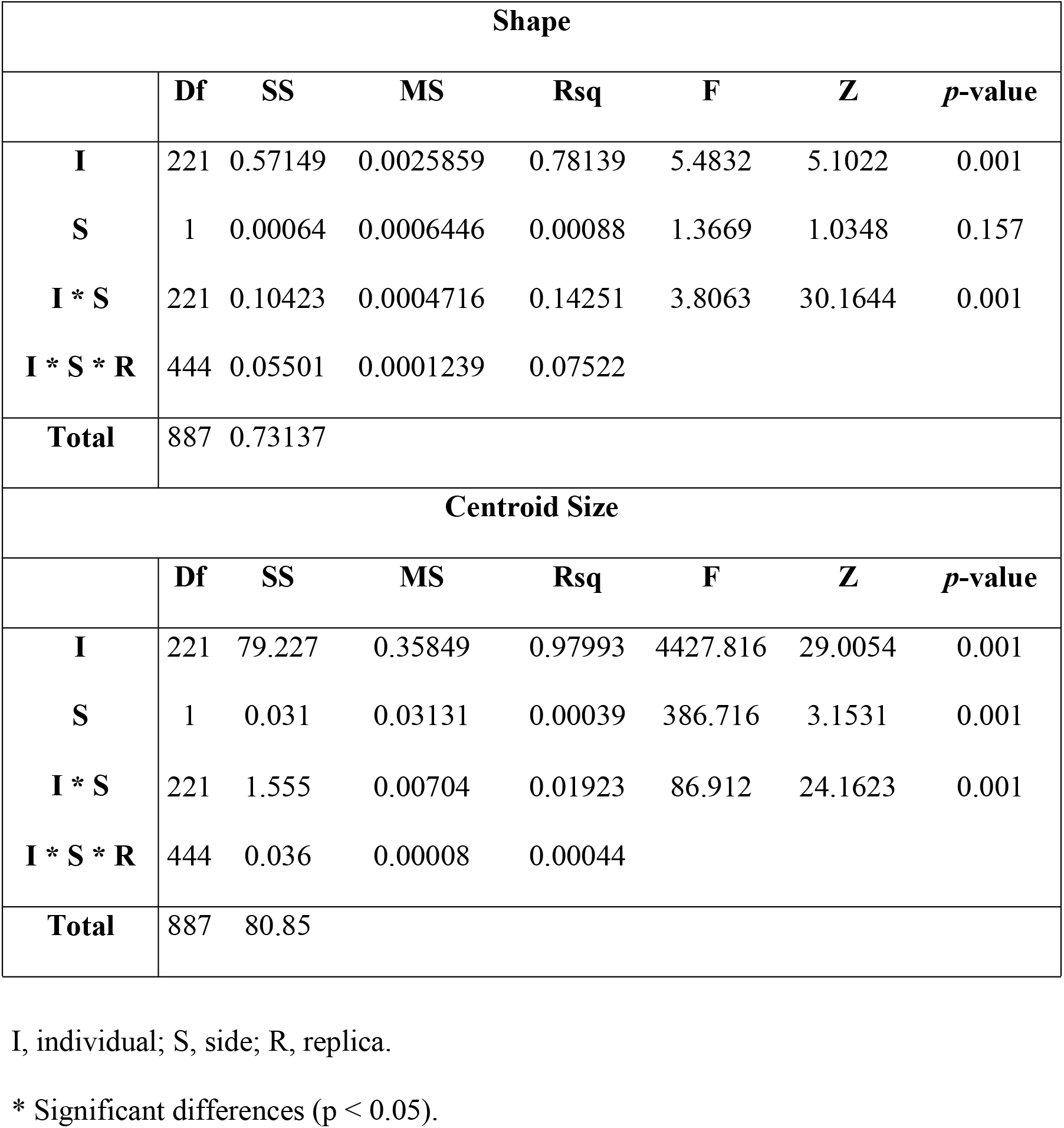
MANOVA of Procrustes for the shape and centroid size of wings in males of *A. aegypti.*

| Shape |  |  |  |  |  |  |  |
| --- | --- | --- | --- | --- | --- | --- | --- |
|  | Df | SS | MS | Rsqr | F | Z | p-value |
| **I** | 221 | 0.57149 | 0.0025859 | 0.78139 | 5.4832 | 5.1022 | 0.001 |
| **S** | 1 | 0.00064 | 0.0006446 | 0.00088 | 1.3669 | 1.0348 | 0.157 |
| **I \* S** | 221 | 0.10423 | 0.0004716 | 0.14251 | 3.8063 | 30.1644 | 0.001 |
| **I \* S \* R** | 444 | 0.05501 | 0.0001239 | 0.07522 |  |  |  |
| **Total** | 887 | 0.73137 |  |  |  |  |  |
| Centroid Size |  |  |  |  |  |  |  |
|  | Df | SS | MS | Rsqr | F | Z | p-value |
| **I** | 221 | 79.227 | 0.35849 | 0.97993 | 4427.816 | 29.0054 | 0.001 |
| **S** | 1 | 0.031 | 0.03131 | 0.00039 | 386.716 | 3.1531 | 0.001 |
| **I \* S** | 221 | 1.555 | 0.00704 | 0.01923 | 86.912 | 24.1623 | 0.001 |
| **I \* S \* R** | 444 | 0.036 | 0.00008 | 0.00044 |  |  |  |
| **Total** | 887 | 80.85 |  |  |  |  |  |

I, individual; S, side; R, replica.
\* Significant differences ( $p < 0.05$ ).

**Table 4.** ANOVA of Procrustes for shape and centroid size on wings of females of *A. aegypti*.

| Shape Anova |  |  |  |  |  |  |  |
| --- | --- | --- | --- | --- | --- | --- | --- |
|  | Df | SS | MS | Rsqr | F | Z | Pr(>F) |
| **I** | 383 | 0.95334 | 0.0024891 | 0.82644 | 7.0429 | 7.8302 | 0.001 |
| **S** | 1 | 0.00045 | 0.0004511 | 0.00039 | 1.2762 | 0.8565 | 0.21 |
| **I \* S** | 383 | 0.13536 | 0.0003534 | 0.11734 | 4.2154 | 30.9891 | 0.001 |
| **I \* S \* R** | 768 | 0.06439 | 8.384E-05 | 0.05582 |  |  |  |
| **Total** | 1535 | 1.15355 |  |  |  |  |  |
| Centroid size Anova |  |  |  |  |  |  |  |
|  | Df | SS | MS | Rsqr | F | Z | Pr(>F) |
| **I** | 383 | 218.322 | 0.57003 | 0.98614 | 5173.998 | 29.9414 | 0.001 |
| **S** | 1 | 0,04 | 0.03959 | 0.00018 | 359.384 | 3.3142 | 0.001 |
| **I \* S** | 383 | 2.944 | 0.00769 | 0.0133 | 69.769 | 26.4014 | 0.001 |
| **I \* S \* R** | 768 | 0.085 | 0.00011 | 0.00038 |  |  |  |
| **Total** | 1535 | 221.39 |  |  |  |  |  |

I, individual; S, side; R, replica.
\* Significant differences ( $p < 0.05$ ).

## 4. Discussion

To our knowledge, this is the first work on an altitudinal gradient in the Andean region identifying differences for wing size and shape of *A. aegypti* males and females in an altitudinal gradient. This is a pattern previously observed in females from *Culex theileri* [18] and *Aedes vexans* [32], West Nile virus and Valley fever vectors, respectively. In *C. theileri*, it was found that wing size and altitude are correlated positively, while in *A. vexans*, these differences were observed for wing size and shape through the altitudinal gradient. Additionally, in *C. theileri* and *A. vexans*, these differences were noted in altitudinal gradients from 808 to 2,130 m (with a difference of 1,322 m) and 808 to 1.620 m (with a difference of 812 m), respectively. Curiously in our case, said difference was observed within an altitudinal range from 1.200 to 1.700 m with a difference of 500 m. For both species, the differences observed regarding wing shape or size could be attributed to variables, like relative humidity and temperature. In turn, in *A. aegypti*, the wing shape and size variation observed may be due to the influence of temperature. In this species, under laboratory conditions, it has been noted that larvae subjected to temperatures between 24 and 35 °C have generated males and females with larger wing size in temperatures from 24 to 25 °C, while at temperatures from 34 to 35 °C, males and females have been obtained with smaller wing size [58]. According to (Atlas, 2019), historical means annual temperatures between 1981 and 2010 for altitudes in our study of 1.200, 1.400, 1,500, and 1.700 m are 20 - 22 °C, 20 - 22 °C, 16 - 20 °C, and 20 – 24 °C, respectively. However, for our case, smaller wings were observed at altitudes between 1,200 and 1,500 m for males, and between 1.500 and 1,700 m for females. Bigger wings were found females at 1.400 m and for males between 1.400 and 1.700 m, which differs from the study already mentioned. Probably, this disparity may be attributed to other variables not measured in this study, like larval density and availability of food. In *A. aegypti* males and females, it was experimentally observed that wing size is correlated negatively in function of the larval density, and positively in function of the availability of food, [59], hence, expecting that with a higher larval density, mosquitoes will have a smaller wing size, and with greater availability of food, there will be bigger individuals. Previously, in *A. aegypti*, it was noted that females with bigger wings have higher survival and greater amount of feeding events through blood [60] and, thereby, increase the probability of transmitting some type of arbovirus [14]. This study did not measure wing length; nonetheless, it has been evidenced that CS has a linear relation with traditional wing length measurements [28], hence, higher CS values would indicate longer wings. In our case, higher CS were found at 1.400 m (Armenia), a city that has historically reported a higher number of dengue cases for the department of Quindío [61].

Furthermore, sexual dimorphism, allometry, and fluctuating asymmetry were found between both sexes for wing size and shape of *A. aegypti*. Sexual dimorphism is a pattern previously observed in morphometric studies for wing shape and size in other Culicidae species [62–65]. Our results suggest allometry between wing size and shape of both sexes, which has been observed in *A. aegypti* [66], as well as in other Culicidae species [27, 67]. Results obtained of fluctuating asymmetry indicate that the wings of both sexes of *A. aegypti* do not have directionality. The fluctuating asymmetry in mosquitoes can be attributed to environmental pressures [68], among them, vector control in urban zones could be one of them, given that previously resistance was detected to organophosphorus compounds, a class of insecticide commonly used for larval control in some of the sampling zones evaluated herein [69].

## 5. Conclusion

In a small scale and in an altitudinal gradient of the Colombian Andes, we found that geometric morphometry permits identifying phenotypic variation for *A. aegypti* wing size and shape. Geometric morphometry studies on wing variation could be used by vector control programs as a diagnostic tool to quantify the dispersion and vector capacity of *A. aegypti.* Future studies must be carried out to test if wing size is related with the vector capacity in this species.

## 6. Acknowledgments

The authors thank office of the vice-president for research, Universidad del Quindío for funding the Project (Grant 828). Gratitude is also expressed to the Biology Program and the Center of Studies and Research on Biodiversity and Biotechnology at Universidad del Quindío (CIBUQ) for providing reagents and equipment, as well as the Health Secretary of Armenia for its support and suggestions during the sampling.

We also thank the members of the Center of Research on Tropical Diseases (CINTROP) for their help and training in identifying the mosquitoes.

This work is dedicated to the memory of the grandmother (La Plita) of Víctor Hugo, who reached the age of 100 years by the time this research ended.

